# Deriving accurate molecular indicators of protein synthesis through Raman-based sparse classification

**DOI:** 10.1101/2021.03.02.433529

**Authors:** N. Pavillon, N. I. Smith

## Abstract

Raman spectroscopy has the ability to retrieve molecular information from live biological samples non-invasively through optical means. Coupled with machine learning, it is possible to use the large amount of information contained in a Raman spectrum to create models that can predict the state of new samples based on statistical analysis from previous measurements. Furthermore, in case of linear models, the separation coefficients can be used to interpret which bands are contributing to the discrimination between experimental conditions, which correspond here to single-cell measurements of macrophages under *in vitro* immune stimulation. We here evaluate a typical linear method using discriminant analysis and PCA, and compare it to regularized logistic regression (Lasso). We find that the use of PCA is not beneficial to the classification performance. Furthermore, the Lasso approach yields sparse separation vectors, since it suppresses spectral coefficients which do not improve classification, making interpretation easier. To further evaluate the approach, we apply the Lasso technique to a well-defined case where protein synthesis is inhibited, and show that the separating features are consistent with RNA accumulation and protein levels depletion. Surprisingly, when Raman features are selected purely in terms of their classification power (Lasso), the selected coefficients are contained in side bands, while typical strong Raman peaks are not present in the discrimination vector. We propose that this occurs because large Raman bands are representative of a wide variety of cellular molecules and are therefore less suited for accurate classification.

## 1 Introduction

Raman spectroscopy is an optical technique that possesses the ability to retrieve highly specific information based on the vibrational modes of the probed molecules. Its non-invasiveness and high specificity make it a technology of choice in various domains that include, for instance, quality control [1] or drug development [2]. Raman is also used in the context of biology and medical applications, but the wide variety of molecular species present in the intracellular environment or tissue, depending on the scale of observation, often makes the measurement less specific. While some applications can exploit resonant responses and achieve sufficient signal to allow ‘classical’ spectroscopy analysis based on band shifts and local intensity changes, such as in the case of blood investigation based on hemoglobin [3, 4] or heme-based compounds [5], most studies have to rely on statistical tools to derive reliable results.

Methods such as principal component analysis (PCA) have been used extensively to analyze Raman data, where the large amount of data points per spectra — typically in the order of a thousand — makes it an ideal candidate for the use of multivariate analysis tools and chemometrics [6]. In the biomedical context, Raman spectroscopy is employed more as a classification tool in conjunction with machine learning algorithms, where the use of supervised learning methods are taking advantage of the high information content of Raman spectra, while compensating for the relatively low specificity of the measurement. Such an approach has been successfully employed for various medical diagnostic applications [7], cellular phenotyping [8], or delineation of diseased tissue in cancer treatment [9, 10]. It is also extensively used in more fundamental research where it can be used to study specific biological processes, including cellular death [11, 12, 13], cellular response [14], infection detection [15] and pathogen identification [16, 17]

On the other hand, Raman spectroscopy has a relatively slow recording rate, which is imposed by the long exposure times required to retrieve with reasonable signal-to-noise ratio the low intensity signals emitted by biomolecules. This implies that it is often challenging to measure large numbers of samples, which are often required in biology to derive relevant findings. In the context of single-cell measurements, recent advances have however demonstrated the ability to measure over thousands of cell samples with this technology [18, 19, 20].

Classification with spectroscopic data based on supervised learning often results in a compromise between the specificity of the model, its stability when applied to new data, and interpretability. In this article, we are focusing on linear methods, which provide very easy to interpret vectors in the mathematical sense, as the model coefficients can be directly understood as representing a given class. However, even in such a simple case, the coefficient distribution within vectors is often complex, making the actual spectroscopic interpretation, which involves relating a given class with underlying molecular species, a challenging task.

We first study the performance of different linear classification approaches, both in terms of performance (accuracy, specificity), and interpretability of the resulting vector. We show in particular how regularized methods yield sparse models that are easier to interpret, and use that approach to study the relevant markers involved during protein synthesis. We apply the algorithms to spectral data acquired from single cells (macrophage-like cell line), where conditions are determined by their immune activation state induced *in vitro*, coupled with drug-induced protein synthesis inhibition. In particular, we study the impact of PCA on the classification characteristics, by comparing the combination of PCA with linear discriminant analysis (LDA) and regularized approaches applied directly to spectral data, namely least absolute shrinkage and selection operator (Lasso).

The feature vectors provided by the Lasso approach are significantly more sparse than the original spectra, since a large portion of the wavenumbers are set to zero by the regularization process that suppresses variables if they do not significantly contribute to classification performance. This makes the separation vector features different from usual Raman data, possibly complicating interpretation. To study how these sparse vectors can be interpreted, we apply Lasso to well-defined conditions, where we induce immune activation, which is known to promote the expression of pro-inflammatory signaling proteins, and compare this condition with a case where protein synthesis is inhibited. Contrary to expectations, the results show that the vectors that provide the most accurate classification do not rely on the main Raman bands characteristic of a cellular spectrum, and instead rely on side-bands and information away from large peaks. This initially counter-intuitive result highlights an interesting aspect of the use of Raman data to classify targets, and we hypothesize that the largest spectral bands are less useful for classification since they are representative of too many molecular species. We also show that the side bands selected for the classification vector are consistent with the known biological effect under study here, namely protein synthesis inhibition.

## 2 Material and Methods

### Cell culture and stimulation

Raw264 (Riken BioResource Center) are cultured in Dulbecco’s modified Eagle medium (DMEM, Nacalai) supplemented with 10% fetal bovine serum (Gibco) and penicillin/streptomycin (Sigma-Aldrich) with 10,000 units and 10 mg/mL diluted at 10 mL/L, respectively. Cells are plated on 10-cm tissue-culture dishes and incubated at 37°C in a humidified atmosphere with 5% CO_2_. For observation in the Raman system, cells are first detached from the dish with a solution containing 0.25% trypsin and 1 mM ethylenediaminetetraacetic acid (Nacalai) for approximately 5 minutes at 37°C. The cell suspension is then plated at a density of 30, 000 cells*/*cm^2^ on quartz dishes (FPI) pre-coated with poly-L-lysine (PLL, Sigma-Aldrich) by immersing the surface in a 0.01% PLL solution (Sigma-Aldrich) for 30 min at room temperature (RT). Cells are then incubated for 5–6 hours to allow them to adhere to the dish substrate. They are then stimulated by replacing the culture medium with fresh DMEM containing lipopolysaccharide (LPS) from E. Coli O111:B4 (Sigma-Aldrich) and/or cycloheximide (CHX, Sigma-Aldrich). Cells are then incubated for 20–21 hours before measurements.

### Raman measurements

The cell culture on quartz dish is washed 2–3 times with phosphate buffer saline (PBS, Nacalai) supplemented with 5 mM of D-glucose and 2 mM of MgCl_2_ (Nacalai) just before measurement with the Raman microscopy system, which has been described previously [21, 22]. Briefly, a 532 nm laser (Verdi, Coherent) is employed as an excitation laser, which is focused onto the sample with a 40× objective (0.75 and 0.95 NA for LPS and CHX experiments, respectively), yielding a power at the sample of 174 and 278 mW/*μ*m^2^, respectively. The back-scattered light is collected by the objective, separated from excitation light by a dichroic and a notch filter (Semrock) before being injected into a 500 mm Czerny-Turner spectrometer (Andor) with a 300 lp/mm grating that spreads the spectral information onto a scientific CMOS detector (Orca 4.0, Hamamatsu) to measure the vibrational spectrum (535–3075 cm^−1^) with an exposure time of 3 s.

Cells are imaged with a quantitative phase imaging (QPI) off-axis digital holography system [23] that is employed to selectively target cells in the field of view. Cells are illuminated with a focused beam that rapidly scans a region covering approximately 30–90% of the cell body that includes both cytosol and nucleus during the exposure for each spectra so as to retrieve a more representative single-cell spectrum, as previously described [24].

### Data processing

Raman spectra are first baseline corrected with cubic spline interpolation, and to account for possible day-to-day variations, data sets from different days are calibrated by interpolating them on a common grid based on a spectrum of pure ethanol measured each day. The silent region (1800–2700 cm^−1^) is then removed, yielding a signal composed of 640 variables out of the original 1024 data points.

All processing is then performed with the *R* program [25] (version 4.0.1). Principal component analysis, linear discriminant analysis and Student’s t-tests are performed with built-in functions. Receiver operating characteristic (ROC) calculations and logistic regression, regularized with Lasso are performed with the *pROC* [26] and *glmnet* [27] packages, respectively. Other calculations are based on scripts developed internally.

When generating a model with Lasso, the regularization parameter *λ* is selected by running 10-fold cross-validation, and using the binomial deviance as a performance metric (see Fig. S1). To further reduce the amount of used variables while ensuring high accuracy, the selected *λ* corresponds to the value that increases deviance by less than one standard deviation compared to the average minimum.

To compare the performance of different models, we employ the cross-entropy (CE), which measures the distance between the expected probabilities derived from the model and the actual ones. The advantage of such a metric compared to the classification accuracy is that it provides the distances to the ideal values, rather than a simple binary indicator, and this produces a higher overall consistency.

## 3 Results and Discussion

In the first part of the article, we study the performance of classification algorithms, in particular by comparing regularized models with standard linear classification methods, and study the influence of employing PCA as a processing step before performing classification.

Classification methods are applied either directly to recorded spectra, or to data first decomposed by PCA to separate the spectral information in orthogonal components, ordered by decreasing importance before applying supervised classification. In particular, we study the combined method PCA/LDA, which has been very popular as a classification tool in vibrational spectroscopy thanks to its relative simplicity and its ability to limit the amount of variables used for classification based on explained variance. This is particularly suitable for low sample sizes, where LDA cannot be employed directly. We compare PCA/LDA with a regularized approach, which limits the amount of variables employed in the statistical model by including a regularization term to reduce the influence of variables that do not significantly contribute to the classification accuracy. In particular, we use the Lasso method, which employs an *L*_1_ regularization term [28] that has the property of reducing the weight of variables to zero unless they are relevant for classification.

### LPS activation induces minute changes in the cellular spectrum

We first study the performance of the different analysis and classification approaches described above by considering the effect of LPS on the Raman spectra of macrophage-like Raw264 cells, which we studied in previous works [14]. Cells stimulated for approximately 20 hours with 100 ng/mL LPS are compared with control conditions. Average spectra are shown in Fig. 1, where only minor differences can be identified by simple inspection. This is expected since the molecular changes occurring upon LPS stimulation are less than, for example, molecular differences between different cell types [19]. The data is composed of measurement sessions spread across 5 days over an interval of approximately 6 months (3 days in December 2017 and 2 days in August 2018). To estimate the performance of the classification algorithms, one day of the dataset is kept aside for use as an independent batch for testing the models. Furthermore, to also assess the long-term stability of the models, another batch of measurements taken around one year later (April 2019) is also used as an additional separate test data set.

**Figure 1:**
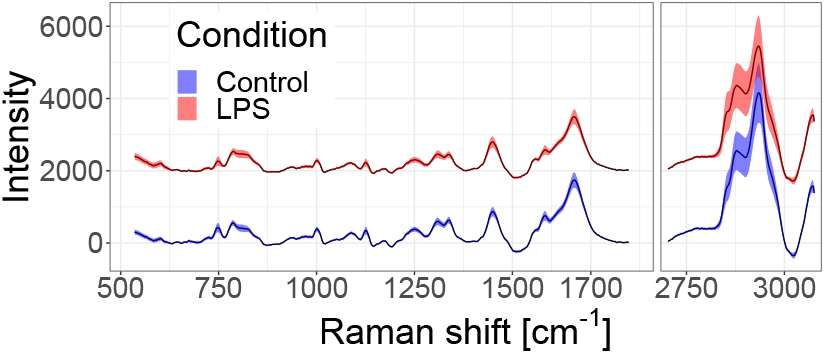
Baseline-corrected average Raman spectra from Raw264 cells, for both control (N=2835) and LPS-exposed (N=2686) cells. Shaded regions represent the standard deviation, LPS spectrum is shown with an offset for visibility.

### PCA/LDA yields lower accuracy and underestimates sample size requirements

Based on the data described above, we created models to classify cells exposed to LPS compared to control conditions, based either on PCA/LDA or the Lasso approach. As PCA/LDA is often used for small sample sizes, we assess the classification performance for increasing training data sizes by computing the resulting CE, as shown in Fig. 2. To account for the variability that can occur due to the choice of subset, the calculations are repeated 10 times with random selection of the training subset.

**Figure 2:**
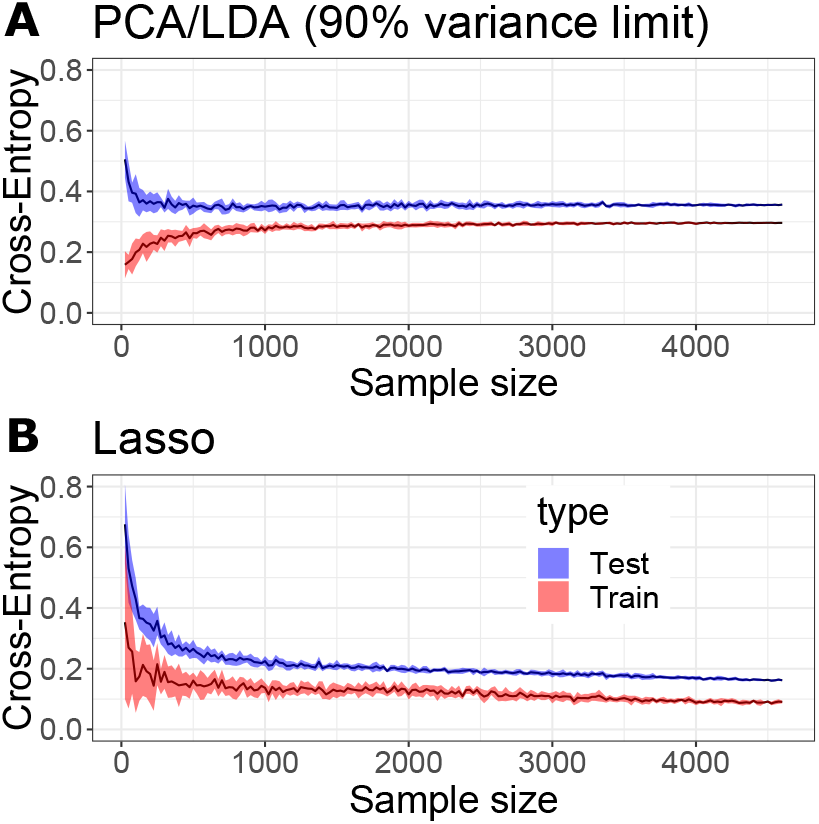
Performance of classification measured by cross-entropy for PCA/LDA with limitation to 90% of explained variance, compared to Lasso with optimization of *λ* at each step with cross-validation. Average of 10 runs with different random selection of subsets, the shaded regions represent the standard deviation.

In the case of PCA/LDA, the amount of variables is limited by including only PCs that explain 90% of the data. This yields a test CE that gradually improves and rapidly reaches a plateau of around 0.355 with a training set size of approximately 600 samples (see Fig. 2A). On the other hand, the Lasso approach (see Fig. 2B) appears to start stabilizing at 0.22 with around 1000 samples, but then continues to improve with increasing training data size, reaching 0.162 at full training size, and it appears that further improvement could be possible. In the Lasso case, the training CE curve shape is unusual as it decreases with data size. This is due to the fact that the regularization parameter *λ* is adjusted at each step, so that the training curve here corresponds to cross-validated performance.

This result demonstrates that PCA/LDA can yield reduced performance in classification compared to Lasso, even when the testing accuracy remains high in both cases (here we obtain 93.9% and 96.0% of testing accuracy, respectively). And, while PCA/LDA performs better here at very small sample sizes, this holds true only for sample sizes below 125, where PCA/LDA has not yet reached optimal performance. Moreover, the evolution of performance with sample size has often been proposed as a metric to determine the required sample sizes for optimum accuracy [29]. As shown in Fig. 2, results derived from PCA/LDA calculation may give the wrong impression that the optimal size has in fact been reached at around 500 samples, while other methods can already perform better by that point, and continue to significantly improve with increasing data sizes. One possible reason for the continuous improvement of Lasso models is that PCA/LDA limited by variance reaches a maximum of around 100 variables and then gradually saturates with sample size, whereas Lasso continues to increase the amount of used variables linearly by adjusting of the variable *λ* (see Fig. S2).

### Inclusion of PCA in models does not contribute to better classification

To study more specifically the influence of using PCA on the prediction models, we next compare the Lasso method either applied directly to the spectral data (as previously), which we refer to as ‘Direct Lasso’, or applied to PCA-decomposed data, denoted as ‘PCA/Lasso’ (in contrast to the use of PCA/LDA in the previous section). As shown in Fig. 3, the performance of both approaches is very comparable, as illustrated by the predicted scores on test data (see Fig. 3A), as well as ROC curves (see Fig. 3B).

**Figure 3:**
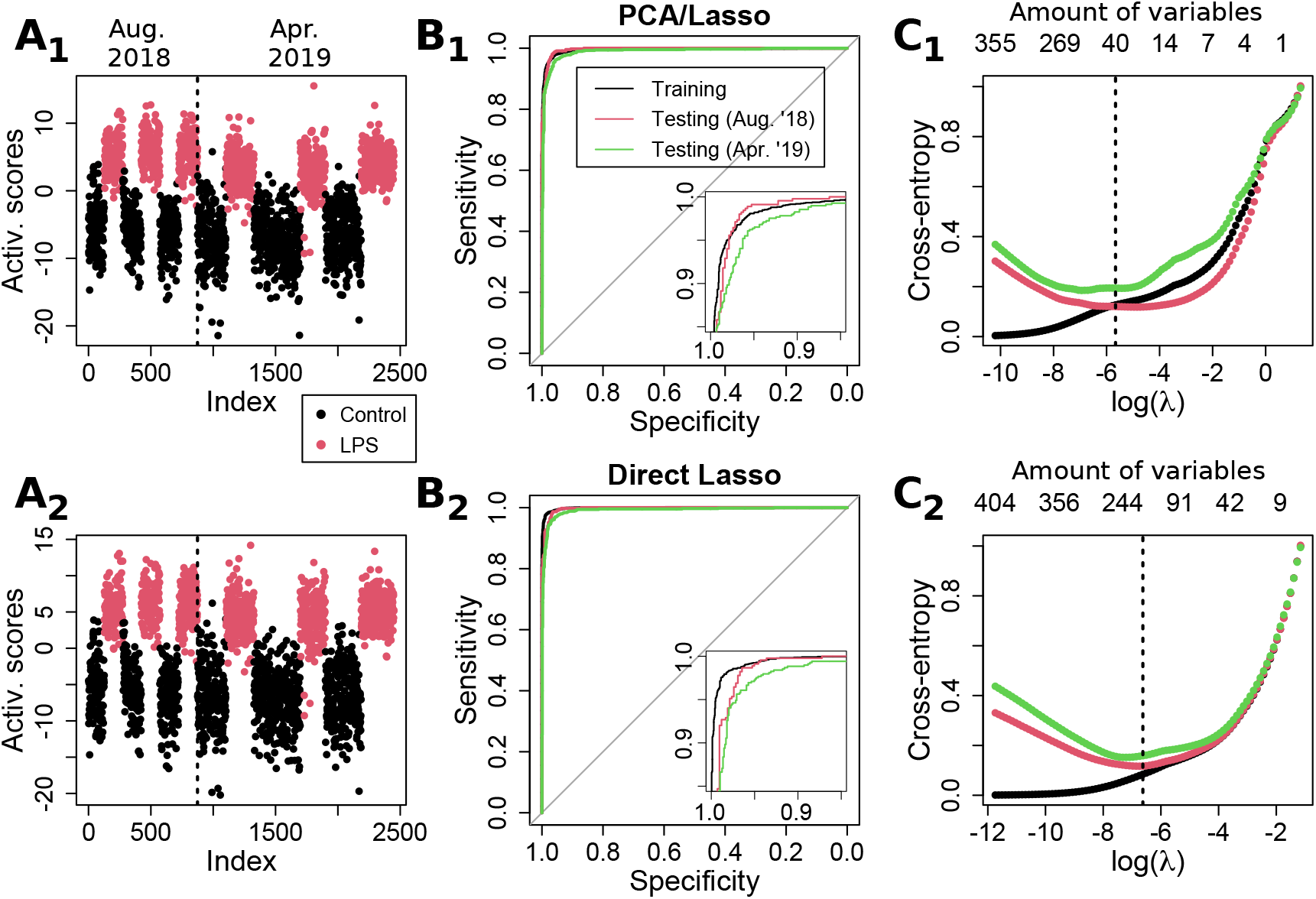
Comparison of performance for Lasso applied either (1) on PCA scores or (2) directly on spectral data. (A) Activation scores for test data (both for a close independent day (Aug. ’18) and data acquired approximately 1 year later (Apr. ’19). (B) ROC results for both methods, showing comparable performances. (C) Cross-entropy as a function of the regularization parameter, showing how the best *λ* changes depending on the type of test data. The optimal *λ* as determined by cross-validation is shown by a dashed line. The corresponding amount of used variables is shown on the top axis.

Some differences can be identified between the two methods when studying the CE as a function of the regularization parameter, as shown in Fig. 3C. While the curves are rather consistent between the two sets of test data in the case of direct Lasso, there is a significant loss of performance for the second set in the case of PCA/Lasso. Furthermore, there is also a shift in the optimal *λ* value, implying that more variables are required to maintain performance, illustrating a reduced stability of the model.

Interestingly, the Lasso average computation time (on 10 runs, 2.9 GHz i7-7820HQ CPU) is shorter when applied to PCA scores (9.5 ± 0.2 seconds) compared to 12.4 ± 0.1 s for direct Lasso. This can be explained by the fact that PCA loading vectors are orthogonal, accelerating the convergence for the Lasso procedure. However, when also taking into account the actual PCA computing time, the PCA/Lasso requires in total 15.4 ± 0.2 s, making it slower overall than direct Lasso.

It can also be seen that the PCA/Lasso approach outperforms PCA/LDA (shown in the first section of results). This can be explained by the fact that Lasso selects PCs within the whole range of variables, while the variance limit implies that only the first 113 out of 640 PCs are used. Interestingly, this shows that the use of high-order coefficients in the case of Lasso does not necessarily create a less stable model. Also a low-order PC is not necessarily providing a strong separation as illustrated by the fact that PCA/Lasso selects only 55 variables out of the 113 variables within the 90% variance limit (see Fig. S3).

These results overall indicate that there is no benefit in employing PCA during the creation of statistical models for prediction. It can even result in some loss of stability, also coupled with an increase in the model complexity, as both PCA loading vectors as well as Lasso coefficients must be used in conjunction to retrieve prediction scores.

Nevertheless, these results overall demonstrate the ability of generating highly stable models that can be employed to predict the immune activation state of individual cells after stimulation purely based on Raman data, despite the high complexity of such cellular changes. It is possible to achieve stability across data taken over a span of at least 8 months, with an independent day within the range of measurements that includes training data, and then with data recorded approximately one year later.

### Direct Lasso provides sparse, less noisy separation vectors

One very valuable feature of classification models based on a linear separation such as LDA or logistic regression is that the resulting coefficients can be used directly; it is possible to retrieve the classification scores by multiplying the coefficients with the input data. This implies that these coefficients can be interpreted as a ‘separation vector’ that indicates which variables distinguish the experimental conditions under study. The vector obtained by PCA/Lasso for LPS stimulation is shown in Fig. 4A, where multiple features can be identified, although the vector is rather noisy due to the inclusion of high-order PCA loading vectors by the classification model.

**Figure 4:**
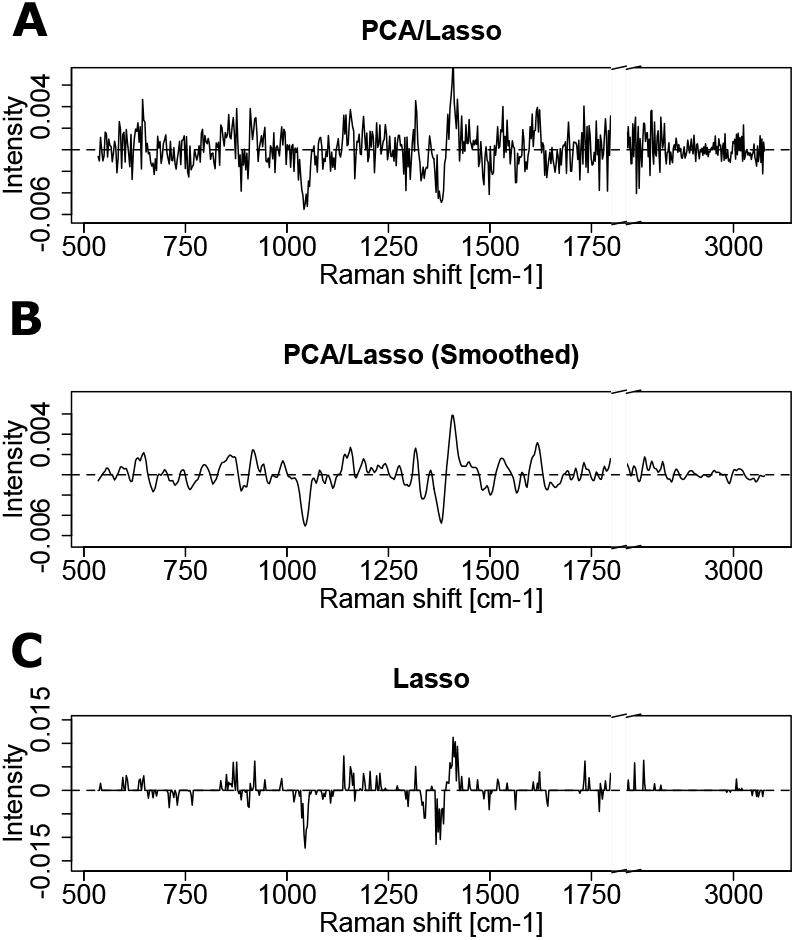
Separation vectors leading to the activation scores displayed in Fig. 3. (A) PCA/Lasso case, where the vector is obtained by combining PCA loading vectors and Lasso coefficients. The noise is due to the inclusion of high-order components. (B) Smoothed version of the PCA/Lasso vector. (C) Sparse vector obtained in the case of direct Lasso.

On the other hand, the vector derived from the direct Lasso model (see Fig. 4C) has sparse features due to the nature of the *L*_1_ regularization, so that all present features have a significant role in the separation, and finer features are easier to identify thanks to the absence of background noise. Nevertheless, the two Lasso and PCA/Lasso vectors share multiple identical features, which are more easily visible by looking at a smoothed version of the PCA/Lasso one (quadratic Savitzky–Golay filter, window size 16, see Fig. 4B), which indicates a certain consistency in the molecular basis of the classification. Furthermore, the most prominent features are also consistent with previously reported results, where PCA/Lasso has been employed to retrieve such vector and interpret the molecular species involved in the case of LPS-exposed cells [14].

One striking point in the features employed for separation is the absence of significant coefficients in the strong regions of the Raman spectrum, such as the C-H stretching region (2870– 3000 cm^−1^) or the strong bands representative of biomolecules in the fingerprint region (CH_2_ interaction, 1420–1480, CC, CO, 1550–1700 cm^−1^). Furthermore, the largest features in the separation vector in Fig. 4C are located outside the main bands and even occur within regions with the smallest intensity. This is unexpected, as it is known that LPS stimulation induces multiple signaling cascades that result in the secretion of pro-inflammatory proteins (cytokines) [30], which could then contribute in such Raman bands.

### Inhibition of protein synthesis yields large spectral changes

As shown above, the direct Lasso method uses statistical analysis and produces a separation feature vector that is sparse, and demonstrates the classification does not rely on the strongest and most common Raman bands. This is then atypical compared to many methods of Raman classification. To better understand the link between the identified separation vector and the underlying molecular differences between cell conditions and induced by biological functions, we also performed experiments within a well-understood model, where we specifically inhibited protein synthesis during cell activation through the application of cycloheximide (CHX), which blocks RNA translation. We stimulated Raw264 cells with 50 ng/mL LPS, and employed simultaneously a concentration of 1 *μ*g/mL CHX. These concentrations ensure that the secretion of IL-6 remains close to baseline levels, while minimizing cytotoxic effects that are known to occur during co-exposure of LPS and CHX [31] (see Fig. S4 for details). The resulting conditions are therefore either Control/LPS to study the cellular immune response (as studied above), or LPS/LPS+CHX to observe the inhibition of pro-inflammatory proteins.

The resulting spectra are shown in Fig. 5A, where the Raman spectra are again very similar between the two conditions. However, with LPS and CHX, very significant changes can be identified when looking at the difference of the average spectra normalized at 2933 cm^−1^ (see Fig. 5B), where an overall decrease in most bands is present, consistent with the blockage of a primary cellular function. These differences are indeed much clearer than the ones occurring purely upon LPS exposure, where most features are significantly smaller, apart from the large difference at 2850 cm^−1^. It can be surprising to find only negative features in the case of CHX blockage, as an accumulation of mRNA could be expected to occur upon inhibition of its translation into proteins, but it is also known that gene expressions can vary under CHX exposure, and that such effects can be pathway-dependent [32]. This in turn can create an imbalance in the secreted cytokines as certain signaling proteins can be released by macrophages without requiring *de novo* protein synthesis [33].

**Figure 5:**
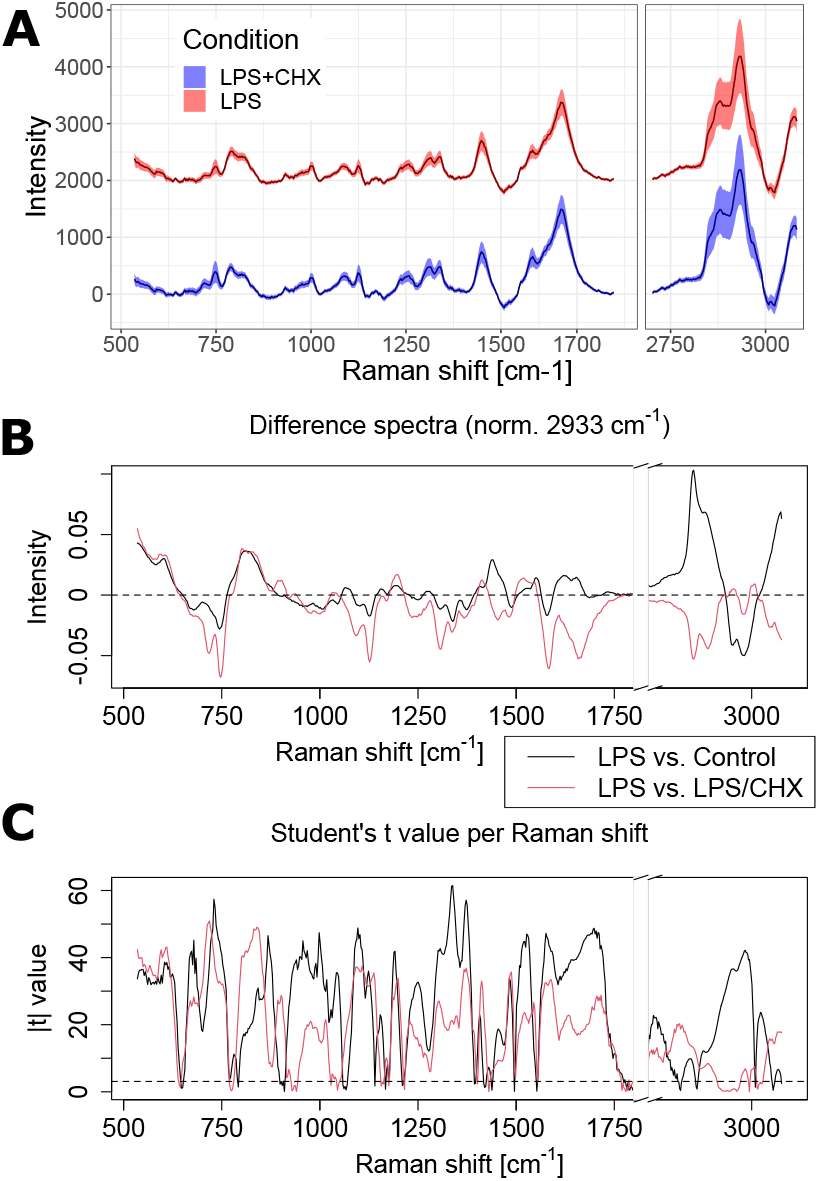
(A) Baseline-corrected average Raman spectra from Raw264 cells, for both LPS (N=2569) and LPS+CHX (N=2512) conditions. Shaded regions represent the standard deviation, LPS spectrum is shown with an offset for visibility. (B–C) Comparison of control/LPS (see Fig. 1) and LPS/LPS+CHX (see Fig. 5A) conditions, shown for the (B) average difference spectra and (C) the absolute value of the two-tail Student’s *t*-test for each Raman shift. The dashed line shows the threshold for *p* < 0.001.

To further understand the contribution of each Raman shift to the separation of classes, we also employ the Student’s *t*-test, applied individually to each wavenumber value. The absolute value of the *t* parameter is displayed in Fig. 5C, for both experimental conditions. It can be seen that while most values are highly significant (the |*t*| value corresponding to *p* < 0.001 is represented by a dashed line), the significance is indeed lower in the C-H stretching region, which can be attributed to the larger variations in this range. Interestingly, LPS vs. control results appear to be more significant, although the classification performance is lower than when blocking protein synthesis, as discussed below. There is also not much correlation between significance and the separation vector displayed in Fig. 4C, as its main features (1045 cm^−1^ negative peak, 1370/1420 cm^−1^ differential shape) are not linked with larger |*t*| values. Overall, these results validate the non-intuitive choice of features outside the main Raman peaks in order to have robust and accurate classification.

### Raman classification vector relies on molecular indicators consistent with known CHX effects

We then generate a statistical model to classify cells exposed to LPS and blocked with CHX, by employing the direct Lasso approach as described previously, and applying the model to one independent day of experiment. The resulting ROC curve is displayed in Fig. 6A, which corresponds to an overall accuracy of 96.8%, slightly higher than in the case of LPS versus control. The resulting separation vector, shown in Fig. 6B, has 122 non-zero values against 162 in the case of LPS vs. control, showing that the model requires less features to reach a higher accuracy, a sign of better stability. As before, there is very little correlation between the vector and the significance of the Raman shifts. This can be explained by the fact that while significance identifies the ability of variables to distinguish the average of both populations, the classification model selects a variable depending on its ability to separate as many individual samples as possible.

**Figure 6:**
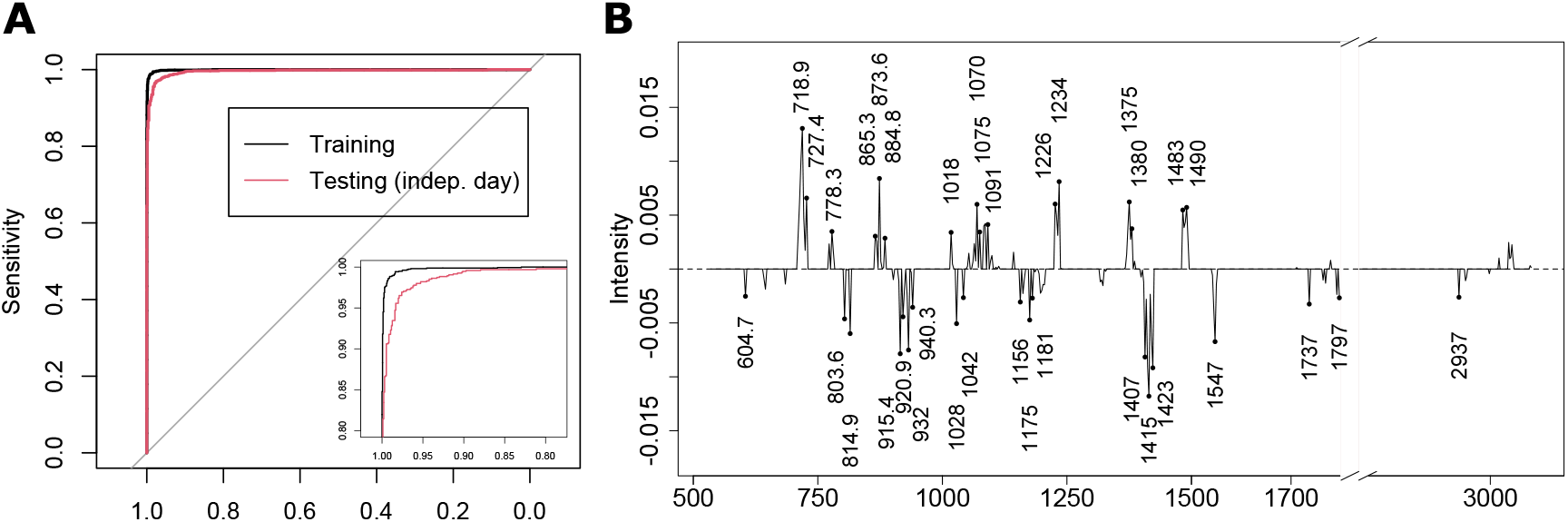
(A) ROC curve for the LPS versus LPS+CHX model. (B) Separation vector corresponding to the model, with the values of the most prominent peaks displayed.

As previously, the selected regions in the separation vector are not located in main region of the spectrum. Furthermore, the most prominent bands that occur in a resonant cellular Raman spectrum (cytochrome c, phenylalanine ring stretching, etc.) are not present here. It is interesting to note that the non-zero regions are essentially contained in groups, despite the correlation that occurs between neighboring wavenumbers, which should contribute to reduce the likelihood of selecting close values under a penalized algorithm. This shows that specific regions in the spectrum are the most powerful to efficiently separate the classes under study. A tentative band assignment is provided in Table S1, where bands are separated by their sign — positive and negative contributions being representative of LPS+CHX and LPS conditions, respectively — and ordered by decreasing strength of the largest value in the band. As it is challenging to assign meaning from sparse regions where other peaks might be present in the original spectrum but not retained in the separation vector, assignments are expressed as possibilities.

It can be seen from this analysis that while there are different possible assignments, most positive bands can be linked to nucleobases such as adenine (720, 1375, 1485 cm^−1^), guanine (1485 cm^−1^) or uracil (780, 1234 cm^−1^), ribose-phosphate (865, 1018 cm^−1^) or DNA/RNA backbone (phosphate, 1091 cm^−1^). This would be consistent with the accumulation of RNA material that occurs upon the blockage of mRNA translation into proteins. On the other hand, the negative bands are less clear in their assignments, which display more variety. Nevertheless, while some weaker bands could be attributed to nucleobases (uracil, 645 cm^−1^ or guanine, 1325 cm^−1^), most bands seem to be related either to protein structure (*α*-helix, 933 cm^−1^), amino-acids (carboxyl groups, 1404 cm^−1^), or amino-acids residues (tryptophan 1548 cm^−1^ or tyrosine/phenylalanine, 1181 cm^−1^), along with DNA/RNA structure (ribose-phosphate, 920 cm^−1^ or A-form helix 815 cm^−1^). This again would be consistent with expected effects of CHX, where its absence is characterized by bands related to the presence of proteins, along with other components possibly due to differences in DNA/RNA constituents.

While it is possible in this case to interpret the LPS/LPS+CHX separation vector thanks to its relative simplicity compared for example with the control/LPS vector in Fig. 4, it remains difficult due to the selectivity of the bands that represent only a fraction of the peaks of a given molecular compound, which is a collateral cost of choosing them strictly based on their classification specificity. On the other hand, a straightforward PCA decomposition also provides a certain degree of separation, as illustrated in Fig. S5, where scores are plotted for the first twelve PCs. These scores are related to loading vectors whose shape is closer to ‘standard’ Raman spectra (see Fig. S6), which might therefore be easier to interpret, although this of course comes at the cost of specificity, as even the PC providing the clearest separation (PC2) reaches only an accuracy of 71.2%. It should be noted that modifications to PCA were also recently proposed to improve separation by accounting for instrument-based biases [34], although this requires some degree of supervision in the otherwise unsupervised PCA.

## 4 Conclusions

While PCA can provide very valuable information in the context of exploratory analysis, we have shown that its use for the purpose of classification based on spectroscopic data is not beneficial. First, classical linear discrimination as performed by PCA/LDA yields less accurate results than other linear methods such as regularized logistic regression by Lasso. The automatic selection of variables provided in this case performs significantly better than a limitation to lower components based on intrinsic data variance, as employed in PCA-based dimensionality reduction approaches. Secondly, it was shown that the use of PCA does not improve performance compared to classification applied directly to the original variables, i.e. wavenumbers. Furthermore, the resulting separation vector, which can be used to predict the state of new samples through direct dot product in case of linear models, is noisier when based on PCA classification compared to the sparse vector obtained otherwise, making the identification of separating features for interpretation harder.

The results were here obtained in the case of changes induced in a homogeneous population of cells through immune activation, which should therefore be relatively subtle compared to cases where different cell types or strains are compared, for instance. Nevertheless, the models derived here were shown to be highly accurate (> 95%) and stable across measurements acquired over a year apart. The immune activation involves multiple complex and concurrent biological processes that make the interpretation of the separation vector difficult. To validate the meaningfulness of the classification, we therefore studied a simpler case where the synthesis of pro-inflammatory proteins was pharmacologically blocked.

While the study of the spectral differences upon protein synthesis inhibition clearly shows an overall reduction of most bands in the average spectrum with very high significance (*p* < 0.001) across all wavenumbers, the separation vector displays features that are outside of the strongest regions in the spectrum. This shows that the classification ability of a Raman shift is not related to the average difference it bears between the studied classes, nor to the statistical significance of this difference. Even in a case where the synthesis of a specific type of molecular compound — here proteins — is blocked, the most prominent bands representative of proteins are not present in the separation vector. One likely explanation is that such bands, whose origins lie in rather common molecular interactions, can be representative of a wide variety of molecules, and thus cannot act as accurate variables for classification.

Nevertheless, an analysis of the bands present in the separation vector provides a view that is consistent with the understanding of the mechanisms involved, where the specificity and relative strength of the bands present in the sparse vector help the interpretation. The inhibition of LPS-induced protein synthesis is represented mostly by bands related to nucleobases and ribose-phosphate complexes, indicative of an excess of RNA, which is consistent with the blockage of mRNA translation induced by cycloheximide. On the other hand, the observation of LPS application alone points towards an excess of proteins, as shown by the presence of bands related to amino-acids residues and protein structure.

The sparse classification vector derived from the Raman spectra provided by Lasso can therefore provide biologically relevant information by highlighting the specific bands that most contribute to the separation of the experimental conditions under study, even if the most classical bands known to appear in spectra of biological molecules are not used in the classification.

## Acknowledgments

This work was funded by the Japan Society for the Promotion of Science (JSPS) through the Funding Program for World-Leading Innovative R&D on Science and Technology (FIRST Program), by the JSPS World Premier International Research Center Initiative Funding Program, and by the JSPS Grants-in-Aid for Early-Career Scientists (KAKENHI Grant Number JP18K14695).

