## Supplementary file (figures and tables) for "Deriving accurate molecular indicators of protein synthesis through Raman-based sparse classification"

N. Pavillon[1], N. I. Smith[1,2]

[1]Biophotonics Laboratory, Immunology Frontier Research Center (IFReC),

[2]Open and Transdisciplinary Research Institute (OTRI),

Osaka University, Yamadaoka 3-1, Suita, 565-0871, Suita, Osaka, Japan

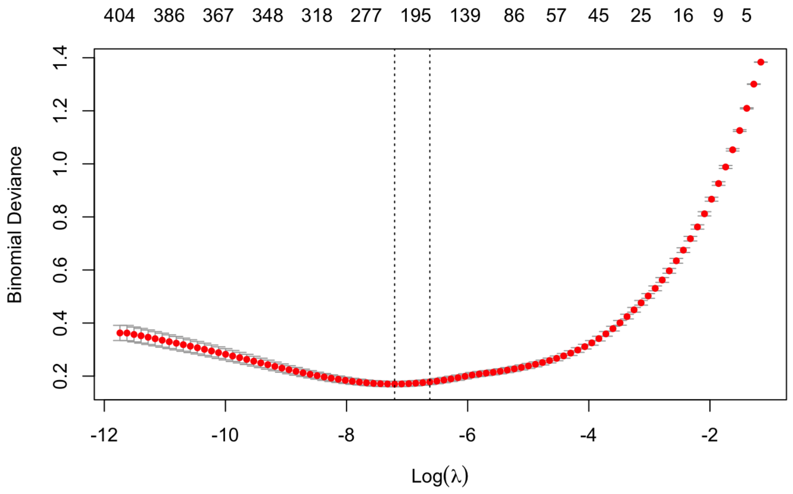


Figure S1: Selection of the regularization parameter $\lambda$ in Lasso through cross-validation, based on binomial deviance. The selected value (right dashed line) corresponds to the $\lambda$ value that increases deviance compared to the average minimum (left dashed line) by less than one standard deviation.

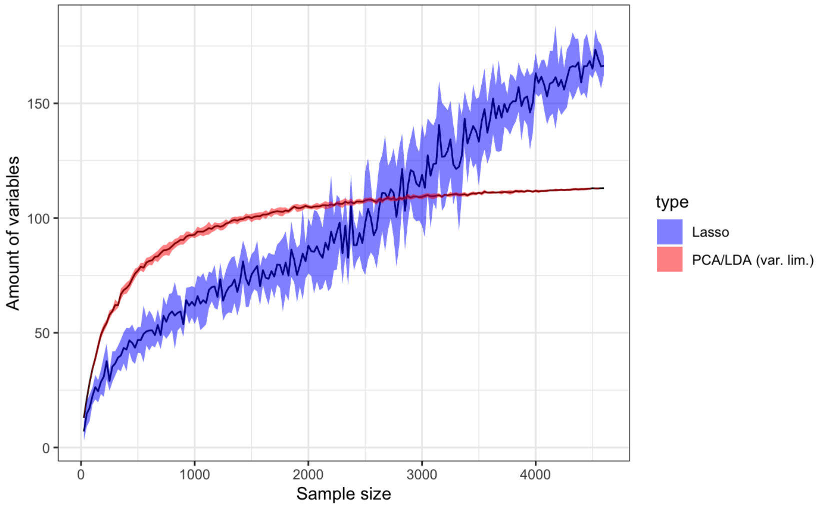


Figure S2: Amount of parameters for Lasso and PCA/LDA in function of sample size. While the limitation to 90% of variance implies that the amount of variables in PCA/LDA rapidly saturates, it keeps increasing in the case of Lasso. The line represents the average of 10 runs with different random selection of training subsets, the shaded regions represent the standard deviation.

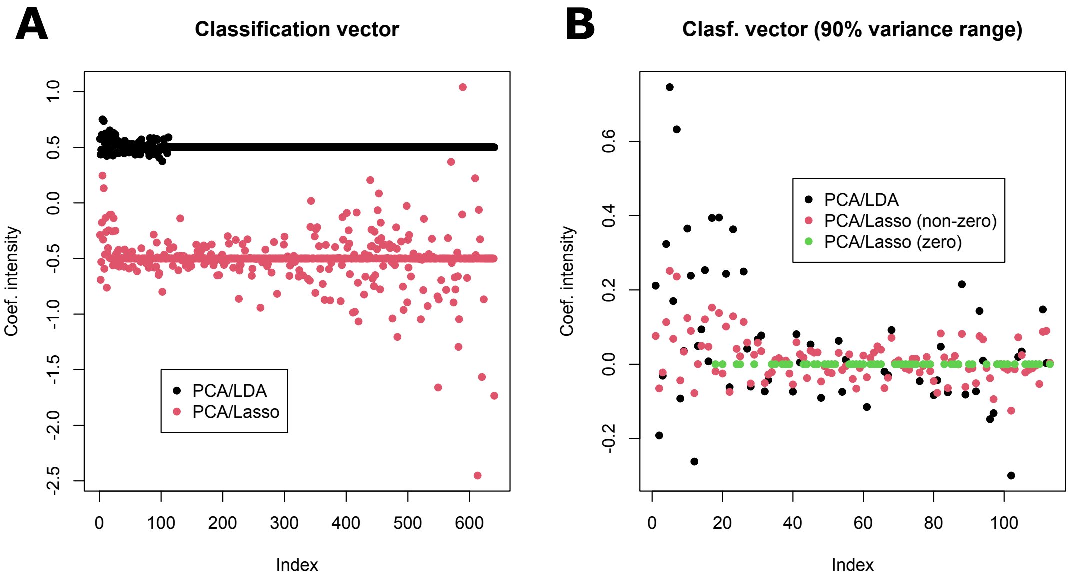


Figure S3: (A) Coefficients of classification models for variance-limited PCA/LDA and PCA/Lasso. The limit on variance implies that selected coefficients are concentrated on low-order PCs, while Lasso selects coefficients in the whole range. An offset of $\pm$ 0.5 has been added for visibility. (B) Coefficients in the low-order PCs within the variance limit, where multiple coefficients are zero (shown in green) in the case of PCA/Lasso.

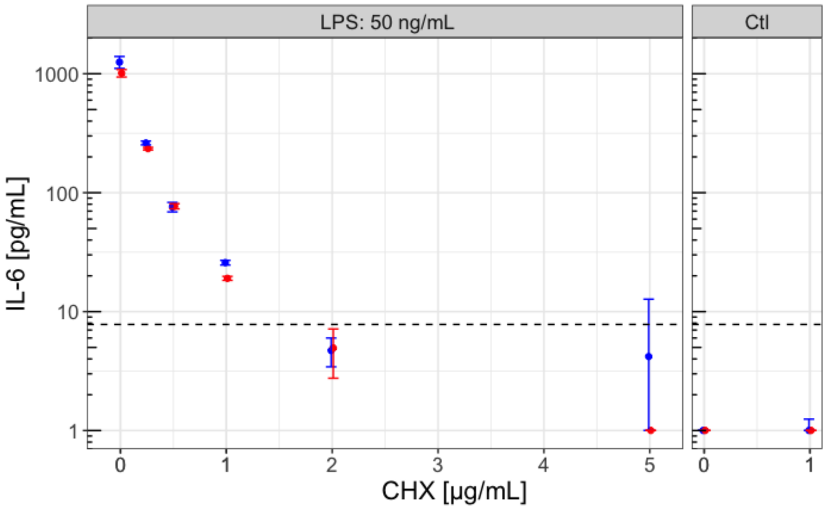


Figure S4: IL-6 secretion measured by enzyme-linked immunosorbent assay (ELISA MAX, Biolegend) in the culture medium of Raw264 cells exposed to LPS (50 ng/mL) and increasing dosage of CHX, represented by the median of triplicate measurements, with the error bar showing the standard deviation. Colors represent 2 independent experiments, and the dashed bar represents the limit of detection of the kit. The manufacturer's protocol was followed, and absorbance was measured at 450 nm.

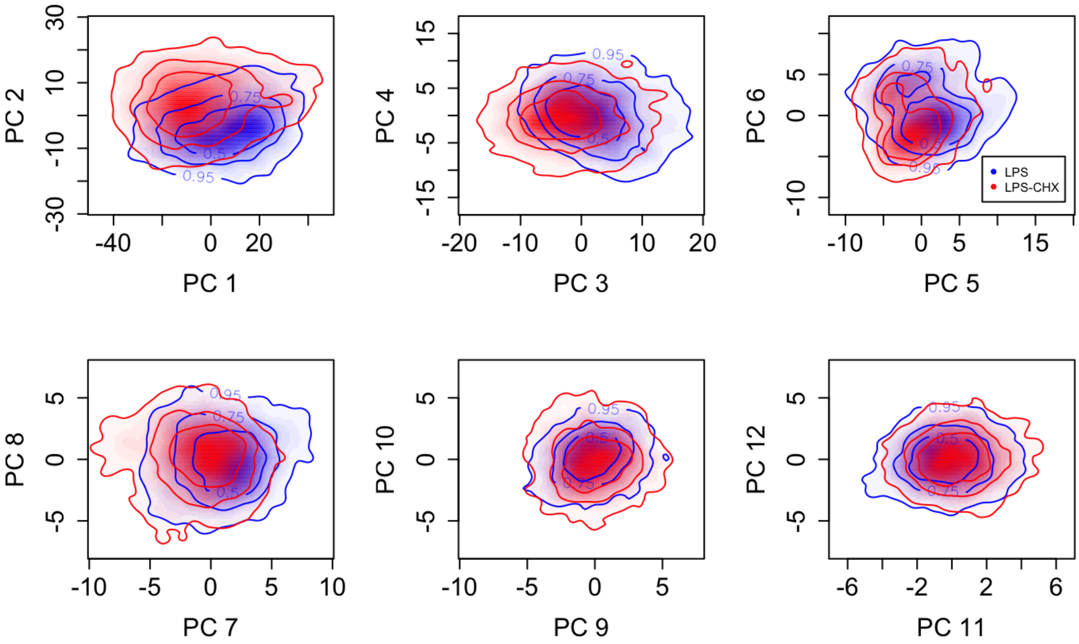


Figure S5: PCA scores for components 1 to 12 on LPS/LPS+CHX data, displayed as density plots. Lines show the limits containing 50%, 75% and 95% of the respective populations.

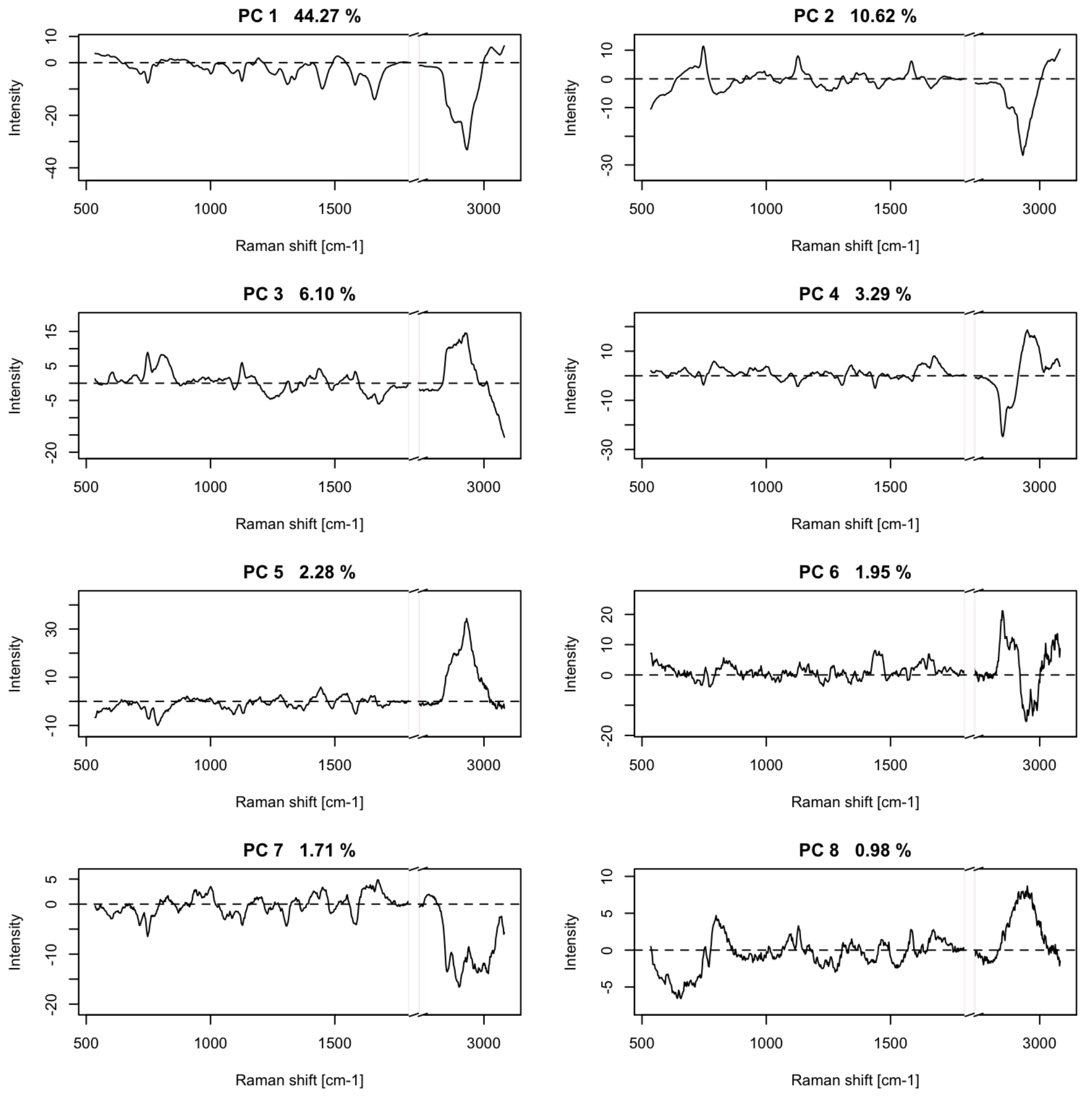


Figure S6: Loading vectors corresponding to scores in Fig. S5. The respective contribution to variance is shown for each vector.

| Band [$cm^{-1}$] | Possible assignment | Vibrational mode | Reported shift [$cm^{-1}$] |
|---|---|---|---|
| | **Positive (LPS+CHX)** | | |
| 710–727 | Adenine | Ring breathing mode [1] | 728 |
| 865–885 | Ribose | Ribose-base linkage [1] | 868 |
| | Tryptophan | Indole ring [2] | 875–880 |
| 1226–1237 | Uracil | Ring stretching [1] | 1234 |
| | Protein secondary structure | Amide III [3], (rand coils [4]) | 1231, 1230–1295 |
| 1050–1113 | DNA/RNA | $PO_2$ stretching | 1092 |
| | Phospholipids | $PO_2$ symmetric stretch [2] | 1080 |
| | Protein primary structure | C-N stretch, | 1066, 1080 |
| | | chain C-C stretch [4] | |
| 1483–1493 | Guanine | CN stretching [1] | 1485 |
| | Guanine, Adenine | Ring mode [5] | 1486 |
| 1367–1386 | Adenine | C-H bending [1] | 1378 |
| | Guanine | Ring stretch [2] | 1370 |
| 773–781 | Uracil, Cytosine | Ring breathing [4] | 782 |
| 1018–1020 | Ribose | CO stretch [1] | 1017 |
| 3037–3046 | | - | |
| | **Negative (LPS)** | | |
| 1399–1423 | Amino-acids | Carboxyl groups side chain [6] | 1404 |
| | Adenine, Guanine | N=C-H bending [1] | 1419 |
| 913–940 | $\alpha$-helix | Backbone CC stretch [7, 4] | 933, 937 |
| | Ribose-phosphate | C-O, C-C stretching [2] | 914–925 |
| 1542–1550 | Tryptophan | Amide II, indole ring [7, 2] | 1551 |
| | - | Amide II, CN stretch [5] | 1548 |
| 801–815 | Lipids | $PO_2$ stretching [4] | 811 |
| | A-form helix | $PO_2$ stretching [2] | 813–816 |
| 1028.5 | - | Coenzyme A [8] | 1027 |
| 1175–1186 | Tyrosine, Phenylalanine | Side chain/Indole ring [7] | 1181 |
| | Tyrosine | CH in plane bend [4] | 1176 |
| 1156–1164 | Ribose | Ribose-phosphate [1] | 1162 |
| | - | Backbone CC, CN [6] | 1060 |
| 1194-1205 | Proline, Tyrosine [8] | - | 1194, 1200 |
| | Pyruvate, Coenzyme A [8] | - | 1197, 1203 |
| 1736.5 | - | CO ester stretch [3] | 1735 |
| 1792–1797 | | - | |
| 1042 | Ribose | Backbone ribose [1] | 1043 |
| 642–645 | Uracil | Ribose-uracil link bending [1] | 647 |
| | Tyrosine | C-C twist [4] | 645 |
| 1317-1328 | Guanine | Ring stretch [2] | 1325 |
| | Amide III | NH, CH stretch [2, 7] | 1300–1340, 1327 |

Table S1: Possible band assignments for the separation vectors of LPS versus LPS+CHX. Regions are listed in the order of decreasing strength.
